# STHELAR, a multi-tissue dataset linking spatial transcriptomics and histology for cell type annotation

**DOI:** 10.1101/2025.07.11.664123

**Authors:** Félicie Giraud-Sauveur, Quentin Blampey, Hakim Benkirane, Arianna Marinello, Paul-Henry Cournède, Stergios Christodoulidis

**Affiliations:** Paris-Saclay University, CentraleSupelec, IHU-National PRecISion Medicine Center in Oncology, Laboratory of Mathematics and Computer Science (MICS), Gif-sur-Yvette, France; IHU PRISM: National Center for Precision Medicine in Oncology, Gustave Roussy, Villejuif, France; Department of Medical Oncology, Gustave Roussy, International Center for Thoracic Cancers, Villejuif, France

## Abstract

Understanding the composition of the tumor microenvironment is critical for cancer research. Spatial transcriptomics profile gene expressions in spatial context, revealing tissue architecture and cellular heterogeneity, but its cost and technical complexity limit adoption. To address this issue, we introduce a pipeline to build STHELAR, a large-scale dataset that integrates spatial transcriptomics with Hematoxylin and Eosin (H&E) whole slide images for cell type annotation. The dataset comprises 31 human Xenium FFPE sections across 16 tissue types, for 22 cancerous and 9 non-cancerous patients. It contains over 11 million cells, each assigned to one of ten curated cell-type categories designed to accommodate a pan-cancer setting. Annotations were derived through Tangram-based alignment to single-cell reference atlases, followed by slide-specific clustering and differential expression analysis. Co-registered H&E images enabled extraction of over 500,000 patches with segmentation and classification masks. Quality control steps assessed segmentation accuracy, filtered out low-confidence regions, and verified annotation integrity. STHELAR provides a reference resource for developing models to predict cell-type annotations directly from histological images.

## Background & Summary

Cancer, a leading cause of mortality worldwide with nearly 10 million deaths in 2020, represents one of the most pressing public health challenges. Defined by uncontrolled cellular proliferation and invasive characteristics, cancer includes a broad spectrum of diseases affecting various tissues and organs ^1^. Modern oncology has evolved to view cancer as a complex ecosystem known as the tumor microenvironment (TME). This evolving environment comprises cancer cells and a variety of non-malignant cell types embedded in a vascularized extracellular matrix. Immune, stromal, and vascular cells within the TME significantly influence tumor progression and resistance to treatments, and the composition of this environment varies considerably depending on tumor location, primary or metastatic sites, histological subtype, molecular features, previous systemic treatment or irradiation, response to treatment, and patient-specific characteristics. Consequently, a detailed characterization of the TME is essential for the development of effective anticancer therapies ^2^.

Spatial transcriptomics (ST) has emerged as a powerful tool for understanding spatial organization and cellular heterogeneity within the TME, offering insights not possible through bulk or even single-cell RNA sequencing. ST offers the unique advantage of providing spatially resolved gene expression profiles, using either imaging-based or sequencing-based approaches. Imaging-based ST relies on fluorescent labeling and signal intensity to map targeted gene expression, whereas sequencing-based methods utilize spatial barcoding coupled with next-generation sequencing to spatially quantify gene expression within tissues. Among these technologies, the Xenium platform from 10x Genomics stands out as an advanced imaging-based method, providing subcellular spatial resolution while enabling exact alignment to whole-slide images (WSIs) ^3^.

Despite its potential, widespread adoption of ST technology is impeded by logistical barriers, including high cost, and complex experimental protocols and data processing. In contrast, WSIs provide a more accessible resource for research by enabling the digitization of histological slides for computational analysis ^4^. A WSI is a high-resolution digital scan of a tissue section, typically stained with Hematoxylin and Eosin (H&E), capturing the entire area of the slide at microscopic detail. While traditional histopathological assessment in clinical settings is still predominantly performed using optical microscopy, the use of WSIs has expanded rapidly in research, driven by advances in whole slide scanning technologies and digital pathology software ^5^. Histological examination of tumor biopsy sections remains a fundamental method in oncology, offering critical insights for diagnosis and prognosis. Recent advances and growing datasets have facilitated the development of sophisticated algorithms capable of processing high-definition WSIs. Specifically, deep learning (DL) models have transformed histological image analysis workflows by achieving remarkable performance in complex clinical applications, including mitosis detection, immune infiltration quantification, cancer subtype classification, tumor grading, and prediction of gene expression profiles from WSIs ^6^.

Among these tasks, cell-level instance segmentation and classification are essential for extracting interpretable and spatially resolved biological information from histological images, and they serve as a foundation for many downstream analyses. The CellViT model, a deep learning architecture based on Vision Transformers (ViT), has recently emerged as a powerful method for automated instance segmentation and annotation of cell nuclei from WSIs ^7^. CellViT was trained and evaluated on the PanNuke dataset, a highly challenging nuclei instance segmentation dataset. This dataset consists of 7,901 H&E patches with nearly 200,000 annotated nuclei across five clinically relevant classes in 19 tissue types ^8^. This model demonstrates significant accuracy in classifying nuclei into classes such as neoplastic, inflammatory, epithelial, dead, and connective cells ^7^.

The rapid progress in histological image analysis has been closely tied to the availability of high-quality annotated datasets. In particular, nucleus-level datasets have played a central role in advancing DL methods for instance segmentation and cell classification. These datasets provide the ground truth necessary to train, validate, and compare computational models at the cellular level. However, achieving a balance between annotation accuracy, cellular and tissue diversity, label granularity, and dataset scale remains a persistent challenge. A number of well-known datasets have significantly contributed to this field, each with distinct advantages and limitations. As mentioned previously, PanNuke ^8^ is one of these reference datasets that offers multi-organ scope, with a publicly accessible license. However its annotations are restricted to coarse phenotype classifications and relatively limited cell counts. Another prominent dataset, CoNSeP ^9^ consists of 41 tiles from 16 colorectal-cancer slides, encompassing approximately 25,000 manually annotated nuclei across seven categories (healthy or malignant epithelial, inflammatory, fibroblast, muscle, endothelial, and other). Although providing meticulous, gold-standard annotations, the CoNSeP dataset is highly limited by its size and single-tissue focus. Similarly, NuCLS ^10^ utilizes crowdsourcing to annotate over 220,000 breast-cancer nuclei into 13 fine-grained categories (e.g., mitotic tumor, plasma cells, neutrophils), including metadata on inter-observer variability. Despite this extensive labeling, NuCLS is restricted by potential annotation noise due to its volunteer-driven approach and is limited to breast tissue. The MoNuSAC 2020 challenge dataset ^11^ comprises around 46,000 nuclei from multiple tissues, validated by expert pathologists from diverse institutions to ensure staining variability. Nonetheless, the dataset excludes biologically significant cell types such as fibroblasts, endothelial cells, and plasma cells. The Lizard dataset ^12^ pushes semi-automatic annotation further, providing classification for nearly 500,000 annotated colorectal nuclei into six classes with detailed immune-specific subtypes. This breadth of immune labels and half-million scale are unique for colon tissue, though reliance on model-assisted annotations risks propagating heuristic bias and all non-colon morphologies are absent. Multi-modal strategies are now emerging, such as NeuLy-IHC ^13^ that registers paired IHC and H&E sections from 19 inflammatory-bowel-disease biopsies, automatically assigning around 200,000 cells to three classes (lymphocyte, neutrophil or other). Despite precise immune annotations, the limited cell-type diversity and tissue pathology constrain its broader application. Immunocto ^14^ aligns multiplex immunofluorescence with H&E in 40 colorectal-cancer slides, annotating over 2.2 million immune cells, but limits cell annotations to immune subsets and groups all other cell types into a single residual category. It is also again restricted to colon tissue. Other notable large-scale segmentation resources include the TCGA nuclei segmentation dataset, which comprises quality-controlled nuclear segmentation masks derived from more than 5,000 whole slide images (WSIs) across 10 cancer types, with approximately five billion segmented nuclei ^15^. However, this dataset does not provide any cell-type annotations. Collectively, these datasets highlight the ongoing challenge of simultaneously achieving high cell counts, diverse tissue representation, and detailed cell annotations, a critical requirement for advancing DL models. The development of such datasets is a necessary step toward improving our understanding of cancer and advancing data-driven pathology.

The present study addresses these limitations by building a pipeline to construct an annotated dataset that leverages high-resolution ST data. The resulting dataset, named STHELAR (Spatial Transcriptomics and H&E histology for Large-scale Annotation Resource), comprises H&E-stained histological image patches with corresponding segmentation masks for nuclei, cell-type classification, and tissue provenance metadata. In addition, the dataset includes per-cell and per-nucleus RNA count matrices, standardized cell type annotations, and clustering results, all organized within fully interoperable SpatialData objects. By leveraging ST data, the method aims to generate a more comprehensive, informative, and large dataset compared to traditional annotation approaches, enabling enhanced cell-type annotation capabilities and potentially expanding coverage to a wider array of cell types. Ultimately, STHELAR could help with the development of models that predict cell types directly from routine H&E histological images. Figure 1 provides a summary of this study. More precisely, the following pipeline was followed to build this dataset:

**Figure 1.**
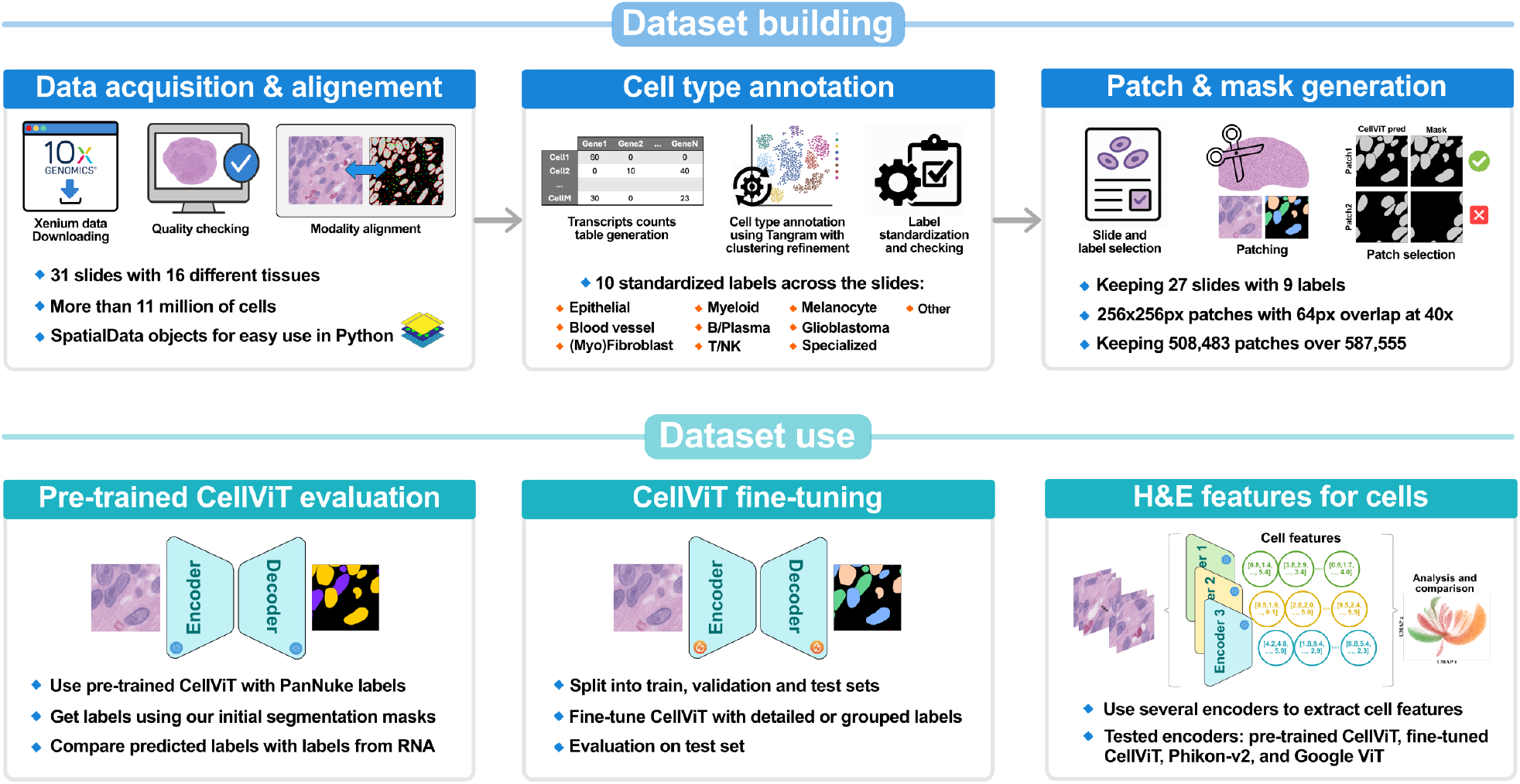
Overview of dataset construction and downstream use case and validations. The top portion outlines the steps of dataset construction. The lower portion illustrates the use case and validation with in particular the fine-tuning of the CellViT model with the generated H&E patches and cell type masks.

### 1 - Building a dataset with 16 different tissues and more than 11 million cells

Data were obtained from the 10X Genomics database ^16^, specifically selecting the Xenium technology that offers single-molecule detection with sub-30 nm localization resolution. This high resolution enables the spatial resolution of RNA expression at the cellular level. H&E slides were aligned with corresponding ST and DAPI data using the Xenium Explorer software ^17^. Detailed characteristics of the dataset are summarized in Table 1.

**Table 1:** Single-cell ST dataset with 31 slides coming from 16 different tissues. The data was obtained from the 10X Genomics website selecting the Xenium technology and only human samples. The nucleus number indicates the total number of segmented nuclei. The gene number corresponds to the number of genes included in the 10X Genomics analysis panel, specifically chosen to enable the identification of major immune cell types and their subtypes. Each slide includes three data modalities: ST, DAPI, and H&E. Gray rows indicate slides that were excluded from the final dataset, along with the reason for exclusion. The “Ref” column links to the corresponding 10x Genomics data reference.

| Slide ID | Tissue | Disease | Nucleus number | Gene number | Ref |
| --- | --- | --- | --- | --- | --- |
| bone_s0 | bone | healthy | 33,801 | 477 | [18] |
| bone_marrow_s0 | bone marrow | acute lymphoid leukemia | 225,906 | 477 | [18] |
| bone_marrow_s1 | bone marrow | healthy | 84,518 | 477 | [18] |
| brain_s0 | brain | brain cancer (glioblastoma multi-forme) | 809,076 | 480 | [19] |
| breast_s0 | breast | breast cancer (ductal carcinoma) | 576,963 | 380 | [20] |
| breast_s1 | breast | breast cancer (ductal carcinoma) | 892,966 | 280 | [21] |
| breast_s2 | same slide than breast_s0 |  |  |  | [22] |
| breast_s3 | breast | breast cancer (lobular carcinoma) | 365,604 | 380 | [22] |
| breast_s4 | same slide than breast_s0 |  |  |  | [23] |
| breast_s5 | same slide than breast_s3 |  |  |  | [23] |
| breast_s6 | breast | breast cancer (II-A, T1cN1MX)) | 692,184 | 5,101 | [24] |
| cervix_s0 | cervix | cervical cancer (III-B,T1bN1MX) | 825,663 | 5,101 | [25] |
| colon_s0 | image not good enough |  |  |  | [26] |
| colon_s1 | colon | colorectal cancer adenocarcinoma | 587,115 | 425 | [26] |
| colon_s2 | colon | colorectal cancer adenocarcinoma | 388,175 | 480 | [27] |
| heart_s0 | heart | healthy | 26,366 | 377 | [28] |
| kidney_s0 | kidney | healthy | 97,560 | 377 | [29] |
| kidney_s1 | kidney | papillary renal cell carcinoma (PRCC) | 56,510 | 377 | [29] |
| liver_s0 | liver | healthy | 239,271 | 377 | [30] |
| liver_s1 | liver | liver cancer | 162,628 | 474 | [30] |
| lung_s0 | alignment not good enough |  |  |  | [31] |
| lung_s1 | lung | lung cancer (NSCLC) | 161,000 | 480 | [32] |
| lung_s2 | same slide than lung_s3 but less relevant cell segmentation |  |  |  | [33] |
| lung_s3 | lung | invasive acinar adenocarcinoma (IB,T2aN0MX) | 275,207 | 5,001 | [33] |
| lymph_node_s0 | lymph node | reactive lymph node (left axilla) | 702,300 | 4,624 | [34] |
| ovary_s0 | ovary | ovary serous carcinoma | 247,636 | 480 | [35] |
| ovary_s1 | ovary | ovarian papillary serous carcinoma (III-B, T3bN0MX) | 401,404 | 5,101 | [36] |
| pancreatic_s0 | pancreatic | pancreatic adenocarcinoma | 190,965 | 474 | [37] |
| pancreatic_s1 | pancreatic | pancreatic ductal adenocarcinoma | 235,099 | 380 | [38] |
| pancreatic_s2 | pancreatic | pancreatic adenocarcinoma | 136,531 | 377 | [39] |
| prostate_s0 | prostate | prostate adenocarcinoma | 184,853 | 5,006 | [40] |
| skin_s0 | image not good enough |  |  |  | [41] |
| skin_s1 | skin | healthy | 90,106 | 377 | [41] |
| skin_s2 | skin | melanoma | 87,499 | 382 | [42] |
| skin_s3 | skin | melanoma | 106,980 | 282 | [43] |
| skin_s4 | skin | primary dermal melanoma | 109,795 | 5,006 | [44] |
| tonsil_s0 | tonsil | reactive follicular hyperplasia | 1,349,620 | 377 | [45] |
| tonsil_s1 | tonsil | follicular lymphoid hyperplasia | 864,388 | 377 | [45] |

The finalized dataset comprises 31 slides derived from 16 distinct tissues, of which 22 slides represent cancerous tissues. In total, the dataset encompasses over 11 million cells prior to patch extraction.

### 2 - Cell type annotation via Tangram with clustering refinement

Accurate cell type annotation is essential, as annotation quality substantially influences the outcomes of subsequent tasks. Several challenges arise when attempting optimal annotation using ST data. A key limitation stems from the inherently lower gene coverage of ST compared to traditional single-cell RNA-sequencing (scRNA-seq), complicating precise cell classification. Another challenge comes from the differences in gene panels, making it difficult to establish uniform, reliable annotations applicable across diverse samples.

Consequently, a slide-by-slide Leiden clustering approach was adopted. To annotate each cluster, we combined differential gene expression analysis and the Tangram method, a deep-learning approach explicitly tailored for ST datasets characterized by limited gene coverage. Tangram generates robust spatial annotations by aligning samples to reference datasets, effectively overcoming resolution and throughput constraints ^46^. Using clustering to refine the labels provided by Tangram resulted in more robust and less noisy annotations.

The final annotations were standardized across slides into the 10 following broad categories, informed by literature ^2^: Epithelials, Blood vessels, Fibroblasts/Myofibroblasts, Myeloid, B/Plasma, T/NK, Melanocytes, Glioblastoma, Specialized, and Other. Table 2 provides a more detailed description of the contents of each cell category. A visual qualitative validation by a pathologist supported the biological relevance of annotations.

**Table 2:** Description of the content of each cell type category.

| Cell type label | Description |
| --- | --- |
| Epithelial | Includes various epithelial cells often specific to individual tissues. For instance, in pancreatic tissue, it includes pancreatic acinar, ductal, and islet cells. |
| Blood vessel | Covers endothelial cells, pericytes, and smooth muscle cells. |
| Fibroblast/Myofibroblast | Contains fibroblasts, myofibroblasts, and mesenchymal stromal cells. |
| Myeloid | Comprises macrophages, monocytes, dendritic cells, neutrophils, mast cells, and plasmacytoid dendritic cells (pDCs). Due to their dual myeloid/lymphoid characteristics and rarity, pDCs were grouped here, with minimal impact expected. |
| B/Plasma | Combines B lymphocytes and plasma cells. |
| T/NK | Includes T lymphocytes and natural killer (NK) cells. |
| Melanocyte | Represents melanocytes or melanoma cells specifically found in skin tissue. |
| Glioblastoma | Identifies glioblastoma cells specific to brain tissue. |
| Specialized | Captures tissue-specific cells such as cardiomyocytes, osteoblasts, osteoclasts, and some endocrine cells. |
| Other | Encompasses cells without marker genes or those with fewer than 10 RNAs. |

Figure 2 shows the distribution of cell types across tissues, and Figure 3 illustrates an example of annotation results at the whole-slide level, shown for the “skin_s2” slide.

**Figure 2.**
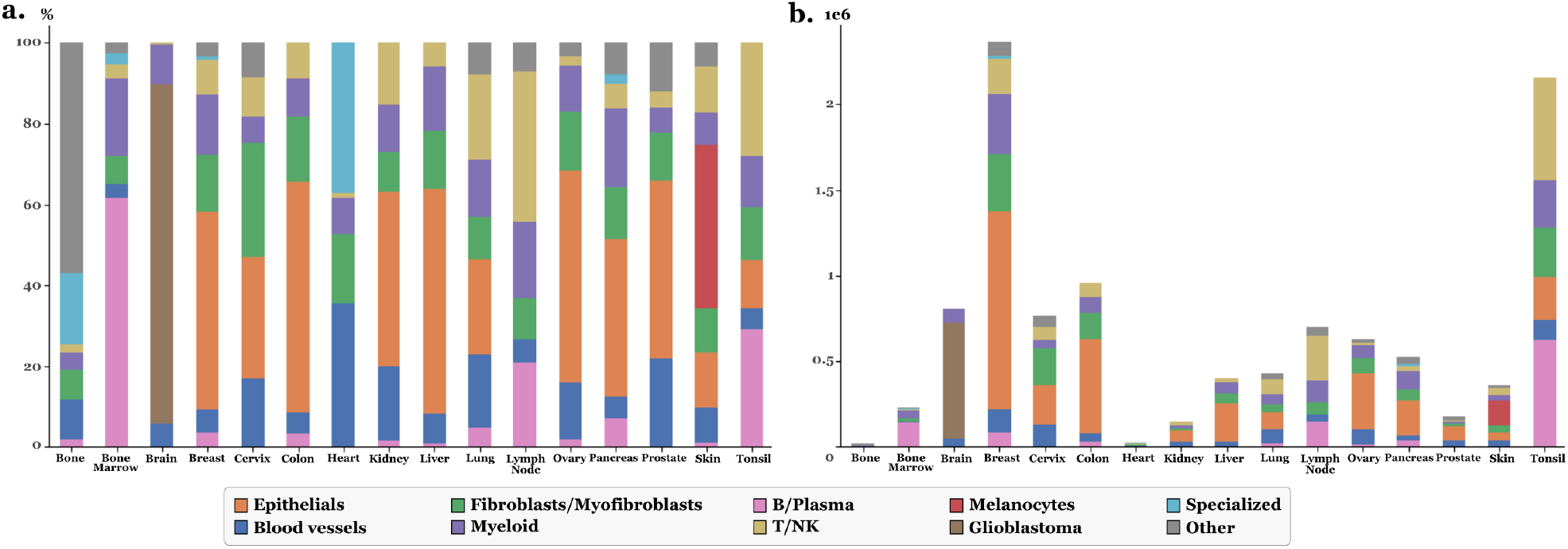
Stackplots illustrating the final dataset for all slides with final cell labels. **a.** Percentage of each cell type per tissue. **b.** Count of cell types in logarithmic scale for each tissue.

**Figure 3.**
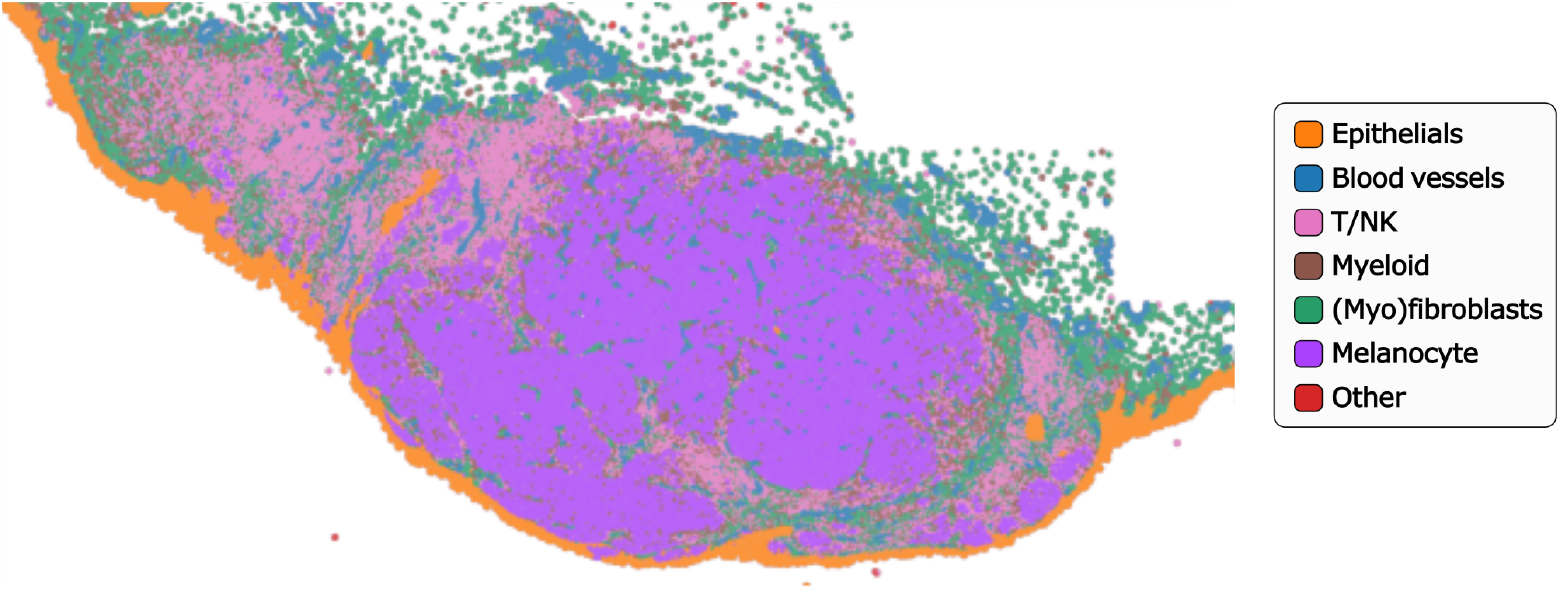
Example of annotation results at the whole-slide level, shown for the “skin_s2” slide.

## 3 - Generating H&E patches with corresponding mask

H&E image patches of size 256 × 256 px with an overlap of 64 px were extracted from each WSI at 40x resolution. Corresponding segmentation and classification masks were generated for each patch in alignment with the PanNuke dataset format ^8^ or with the CellViT model input format ^7^, using the previously established cell-type annotations at the whole cell level. To maintain the generalizability of the dataset, slides with overly specific characteristics (e.g., brain tissue or sample from leukemia patient) were excluded, resulting in a final dataset of 587,555 H&E patches. To support diverse research needs, patches and masks are also provided at a larger field of view with 20x resolution, resulting in a total of 154,814 patches of size 256 × 256 px.

The usefulness and relevance of this dataset was then illustrated by fine-tuning the CellViT model. Details and results are given in the technical validation section, showing its potential to bridge the gap between the rich biological insight offered by ST and the practical limitations of its broader use. By enabling the transfer of ST-derived cell type information to standard H&E images, this dataset facilitates the development of models that approximate ST insights from routine pathology slides.

## Methods

This section describes the procedure for constructing a dataset consisting of H&E image patches with corresponding masks for nucleus segmentation and cell type classification, along with tissue type annotation (i.e., breast, colon, etc.).

### ST data acquisition and processing

Single-cell ST data paired with corresponding H&E images were acquired from the 10x Genomics platform, specifically using datasets generated with Xenium technology that provides both data modalities ^16^. The data selection was restricted to human FFPE tissue sections. Manual quality inspection was performed on each slide using the Xenium Explorer software ^17^ to assess tissue alignment and integrity. The software was also used to generate alignment matrices between the ST data and their associated H&E images. This alignment was achieved by annotating numerous corresponding points across distinct regions on both DAPI and H&E data, enabling the software to compute a transformation matrix. The detailed procedure can be found in the tutorial “Image Alignment in Xenium Explorer” from 10x Genomics ^47^. Slides exhibiting significant misalignment between modalities were excluded from further analysis. Subsequently, each slide was converted into a SpatialData object compatible with Python for integrated analysis of ST data, aligned H&E imagery, and segmentation information. This conversion process used foundational bioinformatics tools provided by the Scverse consortium ^48^, in particular the SpatialData framework ^49^, as well as functionalities available through the Sopa library ^50^, facilitating direct application of the computed alignment matrix within Python. Existing segmentation data, specifically the DAPI-based nuclear segmentation and the default cellular segmentation, were retained along with transcript count tables detailing gene expression at the cell level. The Aggregator tool from the Sopa library was used to determine transcript counts specifically localized to the nucleus for each cell. Since cell identifiers were not consistently preserved throughout data transformations, a shape-based similarity approach for nucleus boundaries was employed to map nuclei accurately to their corresponding cells. This approach ensured the generation of a unified spatial data object encompassing separate transcript count tables for nuclei and entire cells, as well as coherent and unique cell identifiers aligning nuclei and cellular boundaries. The final composition of the dataset is summarized in Table 1.

### Cell type annotation

Cell type annotation posed significant challenges due to the limited number of analyzed genes and differences between gene panels across slides. Multiple approaches were initially explored to overcome these obstacles and achieve robust cell type annotation with consistent and generalizable cell type categories. Early attempts involving batch-corrected clustering across all slides using methods such as Harmony ^51^, scVI embeddings ^52^, and pre-trained embeddings from scGPT, a foundation model trained on extensive single-cell datasets ^53^, failed to yield conclusive results largely due to persistent batch effects. The final choice was to use Tangram annotations combined with cluster-based refinement per slide.

Tangram, a method that learns spatial alignment between sc/snRNA-seq data and a reference spatial dataset, was used for initial cell type annotation ^46^. The selection of an appropriate reference dataset significantly impacts annotation accuracy. Three reference sources were evaluated: CellXGene ^54^, Curated Cancer Cell Atlas ^55^, and the DISCO platform datasets, which provide Deeply Integrated human Single-Cell Omics data organized by tissue ^56^. Quantitative assessments demonstrated that DISCO provided more comprehensive and representative cell types, making it the preferred choice. These Tangram annotations were then refined using a clustering on each slide. Cells with fewer than 10 transcripts were filtered out. Normalization was conducted to equalize transcript counts across cells, followed by log-transformation. Unit variance scaling was applied to each gene, and Principal Component Analysis (PCA) was performed for dimensionality reduction and noise removal. A neighborhood graph was constructed using the first principal components and 10 nearest neighbors. Clustering was conducted using the Leiden algorithm at several resolutions (mainly 0.2, 0.4, and 0.6), with Uniform Manifold Approximation and Projection (UMAP) applied for visualizing cluster distributions alongside Tangram annotations. Marker genes for each cluster were identified using differential gene expression analysis (Wilcoxon signed-rank test with Benjamini-Hochberg correction). Final cell types were assigned by integrating Tangram annotations, UMAP visualizations of marker gene expression across clusters, and literature-based biological context. Figure 4 illustrates the final annotation process using the “ovary s1” slide as an example. Anndata ^57^, Scanpy ^58^, and Rapids-singlecell ^59^ toolkit were used to perform these analyses using GPU acceleration. Final labels were generalized and standardized across slides into the following categories: “Specialized”, “B_Plasma”, “Blood_vessel”, “Epithelial”, “Fibroblast_Myofibroblast”, “Myeloid”, “T_NK”, “Melanocyte”, “Glioblastoma”, “Stem like” (potential stem cells, no clear marker genes), and “Unknown” (cells lacking specific gene markers). This categorization aimed to balance annotation accuracy, biological relevance, and consistency across tissue types (Table 2).

**Figure 4.**
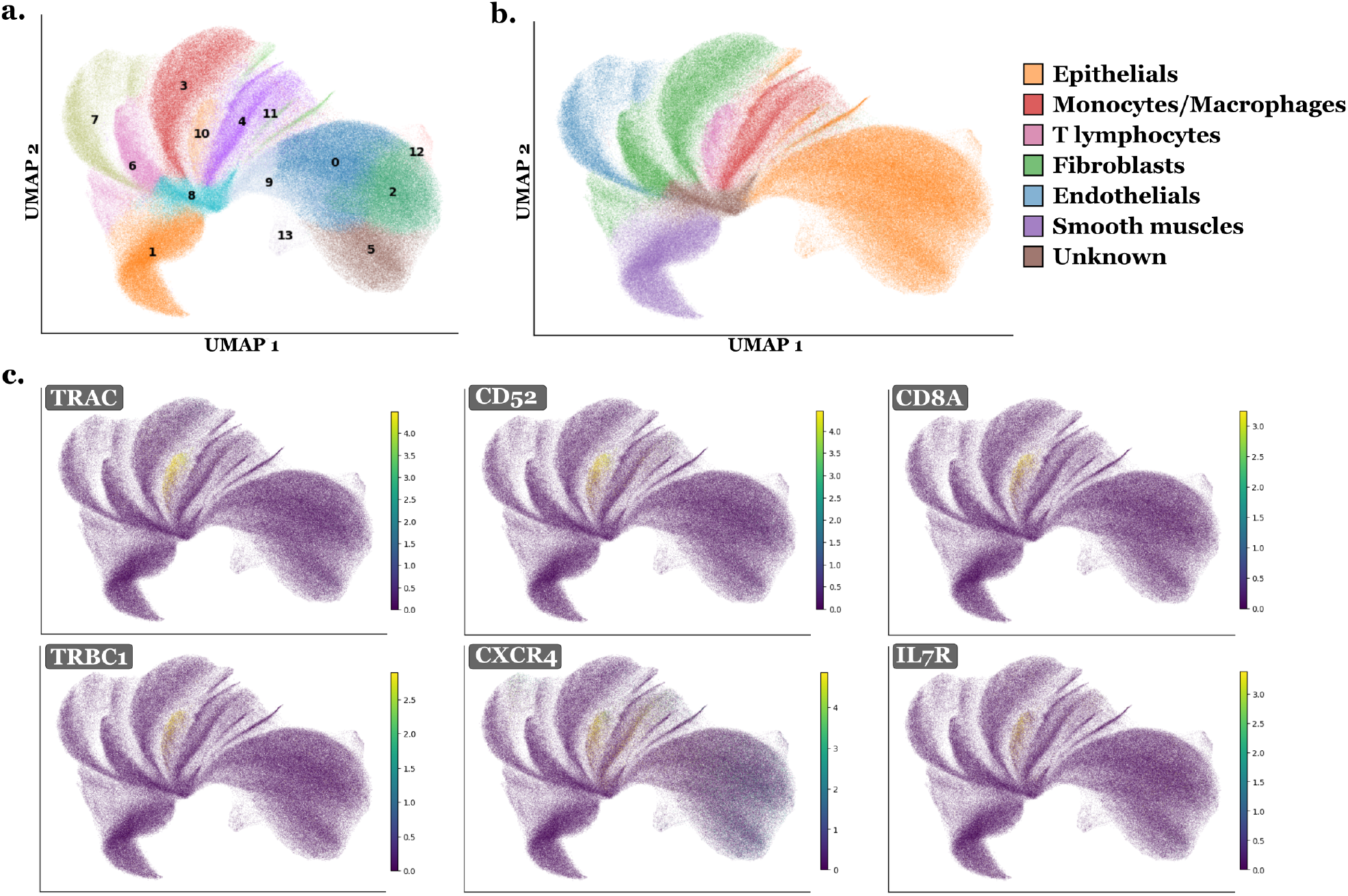
UMAPs illustrating the cell type annotation process. This is an example for the “ovary s1” slide with Leiden clustering at resolution 0.4. In **a.** cells are labeled by their cluster number. Differential expression analysis is performed on these clusters and marker genes are analyzed for each, as illustrated in **c.** with examples of marker genes for cluster 10. In **b.**, cells are labeled with their final labels for the slide, combining information from both marker genes and Tangram labels.

One limitation encountered during annotation was differentiating normal from cancerous cells, as RNA analysis alone did not yield clear distinctions, especially due to the continuous nature of gene expression profiles rather than discrete differences between normal and cancerous cells. Several approaches were explored, including pathway analyses such as KEGG pathway ^60^, which offers a collection of manually drawn pathway maps representing the knowledge of the molecular interaction, reaction, and relation networks for different types of cancers. However, this yielded inconclusive results, and annotations for these categories were retained but marked with limited confidence and not used afterward. A similar challenge arose when attempting to identify dead cells. While low RNA content could serve as a proxy indicator for cell death, it does not conclusively confirm the status. However, qualitative assessment by the pathologist supported this interpretation, as many low-RNA cells exhibited morphological features consistent with cell death. As a provisional measure, these low-RNA cells were grouped within the “Other” category, alongside cells previously categorized as “Unknown”. Both tumor and dead cells represent highly heterogeneous categories, making them particularly difficult to annotate consistently and challenging to predict.

### Generating H&E patches with corresponding mask

H&E image patches of size 256 × 256 px with an overlap of 64 px were extracted from each WSI at 40x resolution, conforming to the specifications required by the CellViT model ^7^. Corresponding segmentation and classification masks were generated for each patch using the previously established cell-type annotations at the whole cell level. To avoid incorporating excessively specialized tissues, such as those containing unique tumor cell populations or distinctive tissue architectures, the slides named “brain_s0”, “bone_marrow_s0”, “bone_marrow_s1”, and “bone_s0” were excluded at this stage. Additionally, the labels “Stem_like” and “Unknown” were consolidated into a single category named “Other”. Thus final mask categories includes: “T_NK”, “B_Plasma”, “Myeloid”, “Blood_vessel”, “Fibroblast_Myofibroblast”, “Epithelial”, “Specialized”, “Melanocyte”, and “Other”.

## Data Records

This work presents a large-scale cell-type annotated dataset derived from 10x Genomics Xenium slides. It includes 31 WSIs with 16 tissue types, each processed into a comprehensive multi-modal object. For each slide, we provide per-cell and per-nucleus transcript count tables with clustering results and standardized cell type annotations, scVI-derived embeddings, cell feature vectors from multiple H&E encoders, nuclei and cell segmentation boundaries, co-registered DAPI and H&E images, and patch coordinates with segmentation metrics. These components are included in a single SpatialData object per slide, with unique identifiers per cell and nucleus to facilitate integration, visualization, and downstream analysis using the Scverse ecosystem ^48^. In total, the dataset contains over 11 million individual cells before patch extraction. The dataset also includes 587,555 H&E image patches with segmentation and cell type classification masks. The overall analyses indicate a good level of confidence in the generated annotations.

This section describes the available data, with a visual summary provided in Figure 5. The dataset ^61^ is stored in the BioStudies database ^62^ and can be accessed online at https://www.ebi.ac.uk/biostudies/bioimages/studies/S-BIAD2146. The cell and patch IDs are kept consistent across all data. To support reproducibility, data related to our technical validation experiments is also provided. For example, the sdata tables (“features_cellvit”, “features_phikonv2”, “features_vit_google”, and “table_scvi”), the PanNuke labels, and the CellViT fine-tuning files and results all relate to content described in the Technical Validation section below.

**Figure 5.**
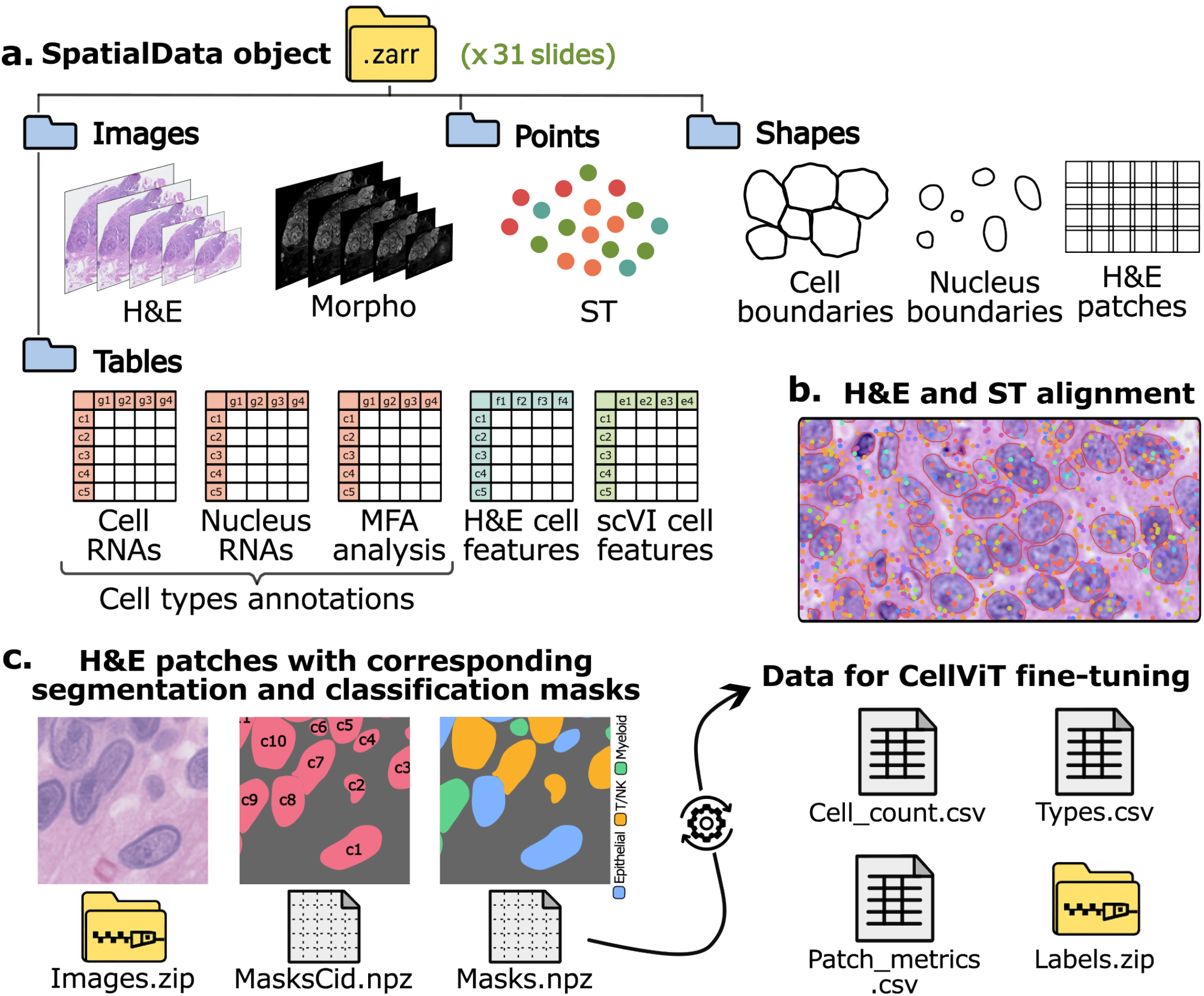
Summary of the dataset. **a.** For each slide, a SpatialData object is provided, including: **H&E**: the H&E-stained whole slide image at multiple resolutions, **Morpho**: DAPI-stained images for nuclear morphology at multiple levels, **ST**: spatial transcriptomics data showing RNA molecule positions, **Cell/Nucleus boundaries**: cell and nucleus segmentation masks, **H&E patches**: coordinates for 40x H&E patches of 256 × 256 pixels with 64-pixel overlap, **Cell/Nucleus RNAs**: RNA count tables per cell and nucleus, along with cell type annotations and clustering labels (“table_cells” and “table_nuclei”), **MFA analysis**: results of a multi-factor analysis (MFA) integrating cytoplasmic and nuclear RNA counts (“table_combined”), **H&E cell features**: extracted H&E features from multiple encoder models (“features_cellvit”, “features_phikonv2”, and “features_vit_google”), and **scVI cell features**: cell-level embeddings from scVI (“table_scvi”). **b.** Example illustrating the alignment between the H&E and ST modalities. **c.** Additional files include: **Images.zip**: a ZIP archive containing all H&E patches at 40x resolution (“images.zip”), **Masks.npz**: per-slide files with cell type masks matching the PanNuke format (“masks_slide-id.npz”), and **MasksCid.npz**: cell ID segmentation masks (“masks_cell_ids_nuclei.zip”). Data formatted for use with CellViT are also provided, including: **Cell_count.csv**: nucleus count per cell type per patch ID, **Types.csv**: tissue type for each patch, **Patch_metrics.csv**: comparison between Xenium based segmentation mask and pre-trained CellViT segmentation prediction, and **Labels.zip**: segmentation and classification masks for all the slide matching the input CellViT format. For model fine-tuning section, 20x resolution images and their corresponding masks and metrics are also included.

### 1. A SpatialData object for each slide in .zarr format

In the **“sdata_slides”** folder, a SpatialData object named **“sdata_slide-id.zarr.zip”** is provided for each slide with all the information and analyses. The SpatialData format can be manipulated using all the tools in the Scverse ecosystem ^48^, making it easy and convenient to use. This SpatialData object includes the following components:

- **“he” image**: H&E slide at several levels (with level 0 corresponding to 40x resolution and level 1 corresponding to 20x resolution), aligned with ST using the Xenium software from 10x Genomics.
- **“morpho” image**: DAPI image at several levels showing each nucleus.
- **“st” points**: Transcripts location on the slide.
- **“cell_boundaries” shapes**: Polygons for cell boundaries with corresponding cell ID.
- **“nucleus_boundaries” shapes**: Polygons for nucleus boundaries with same corresponding cell ID.
- **“he_patches” shapes**: Coordinates for H&E patches of size 256 px with 64 x overlap at 40x resolution, with metrics (Dice coefficient, Jaccard index, and Panoptic Quality) that compares the given mask with pre-trained CellViT prediction.
- **“table_cells” table**: Table that gives the RNA count for each cell, along with preprocessing layer, cell type annotation, cell ID, and clustering.
- **“table_nuclei” table**: Table that gives the RNA count for each nucleus, along with preprocessing layer, cell type annotation, cell ID, clustering, and PanNuke label.
- **“table_combined” table**: Table for MFA analysis with RNA count for cytoplasm and for nucleus, along with clustering analysis and cell type annotations.
- **“features_cellvit” table**: Table that gives the nucleus embeddings for each cell using the pre-trained CellViT model as encoder.
- **“features_phikonv2” table**: Table that gives the cell embeddings for each cell using the Phikon-v2 model as encoder.
- **“features_vit_google” table**: Table that gives the cell embeddings for each cell using the ViT based model from Google as encoder.
- **“table_scvi” table**: Table that gives the scVI embeddings for each cell, using scVI model specifically trained for each slide.

### 2. H&E patches with corresponding segmentation and classification masks

In the **“data 40x/data”** folder, we provide:

- **“images.zip”**: H&E patches for all the slides together, using slide-id patch-id identification.
- **“masks_slides”** folder: Contains a **“masks_slide-id.npz”** file for each slide containing segmentation and cell type classification masks, formatted identically to those in PanNuke.
- **“annot_dicts_ct_1”** folder: Contains the **“label2cat.json”** dictionary which maps the final cell labels in sdata to categories in the classification masks, the **“cat2color.json”** dictionary which maps each category in the masks to a plotting color for visual inspection, and the **“cat2idx.json”** dictionary which maps each category to its corresponding index position within the mask.
- **“masks_cell_ids_nuclei.zip”**: Segmentation masks for all the slides together, assigning to each pixel the cell ID corresponding to its nucleus.
- **“pretrained_CellViT_mask_preds”** folder: Contains a **“instance_map_predictions_slide-id.h5”** file for each slide with the predicted segmentation for each patch using the pre-trained CellViT model.

### 3. Data for CellViT fine-tuning

For the global dataset, the following element can be found in the **“data 40x/data”** folder:

- **“cell_count.csv”**: Table summarizing, for each patch ID across all slides, the nucleus count per cell type. It follows the same format as the cell count.csv file provided by the CellViT authors.
- **“labels.zip”**: Segmentation and classification masks for all slides. Unlike the previous masks.npz files, this format follows the CellViT convention, which separates the original mask into an instance mask and a cell type mask.
- **“patch_metrics.csv”**: Table presenting the evaluation metrics for each mask, using Dice coefficient, Jaccard index, and Panoptic Quality, and comparing with the predicted masks from the pre-trained CellViT model.
- **“types.csv”**: Table giving the tissue type for each patch.

Two fine-tuning have been done with detailed or grouped cell type classes. We provide for each fine-tuning in **“data_40x/finetuning_CellViT_detailed”** or **“data_40x/finetuning_CellViT_grouped”** folders:

- **“cell_count_train.csv”, “cell_count_valid.csv”**, and **“cell_count_test.csv”**: Split versions of the cell co-unt file indicating which patch belongs to the train, validation, or test sets.
- **“dataset_config.yaml”** and **“weight_config.yaml”**: Configuration files required for fine-tuning CellViT.

### 4. CellViT fine-tuning outputs

In each fine-tuning directory (**“data_40x/finetuning_CellViT_detailed”** or **“data_40x/finetuning_CellViT_grouped”**), the subfolder **“output”** contains all results from the fine-tuning and evaluation processes:

- **“config.yaml”**: Configuration file used for CellViT fine-tuning.
- **“logs.log.1–5”**: Log files for fine-tuning.
- **“checkpoint_n.pth”**: Checkpoint for the best model with n the corresponding epoch number.
- **“inference_results.json”**: File containing inference results on the test set.
- **“inference.log”**: Log file for inference.

### 5. Analysis of the fine-tuned CellViT models

For each fine-tuning, the resulting model was applied to all slides. Results can be found in the **“data_40x/finetuning_CellViT_detailed/finetuned_model_analysis”** or **“data_40x/finetuning_CellViT_grouped/finetuned_model_analysis”** folders, that contain a subfolder for each slide ID with the following files:

- **“config.yaml”**: Configuration from the fine-tuning.
- **“inference_results.json”**: Prediction results.
- **“inference.log”**: Log file for inference.
- **“inference_instance_map_predictions.h5”**: Predicted segmentation mask for each patch for all the slides.
- **“cell_features_cellvit.npy”**: Nucleus embeddings for each cell using the given fine-tuned CellViT model as encoder.
- **“pixel_class_gt_mask.pth”**: The pixel count for each cell type category for each nucleus using Xenium based segmentation mask instead of the predicted one.

All analyses were conducted on H&E patches at 40x resolution. However, “images.zip”, “labels.zip”, “masks_cell_ids_nuclei.zip”, “instance_map_predictions_all_slides.h5” (segmentation prediction using pre-trained CellViT), “patch_metrics.csv”, “cell_count.csv”, and “types.csv” are also provided at 20x resolution in **“data_20x/data”** folder for research purposes. For each tissue type, an example patch at 20x resolution along with its corresponding classification mask is shown in Figure 6.

**Figure 6.**
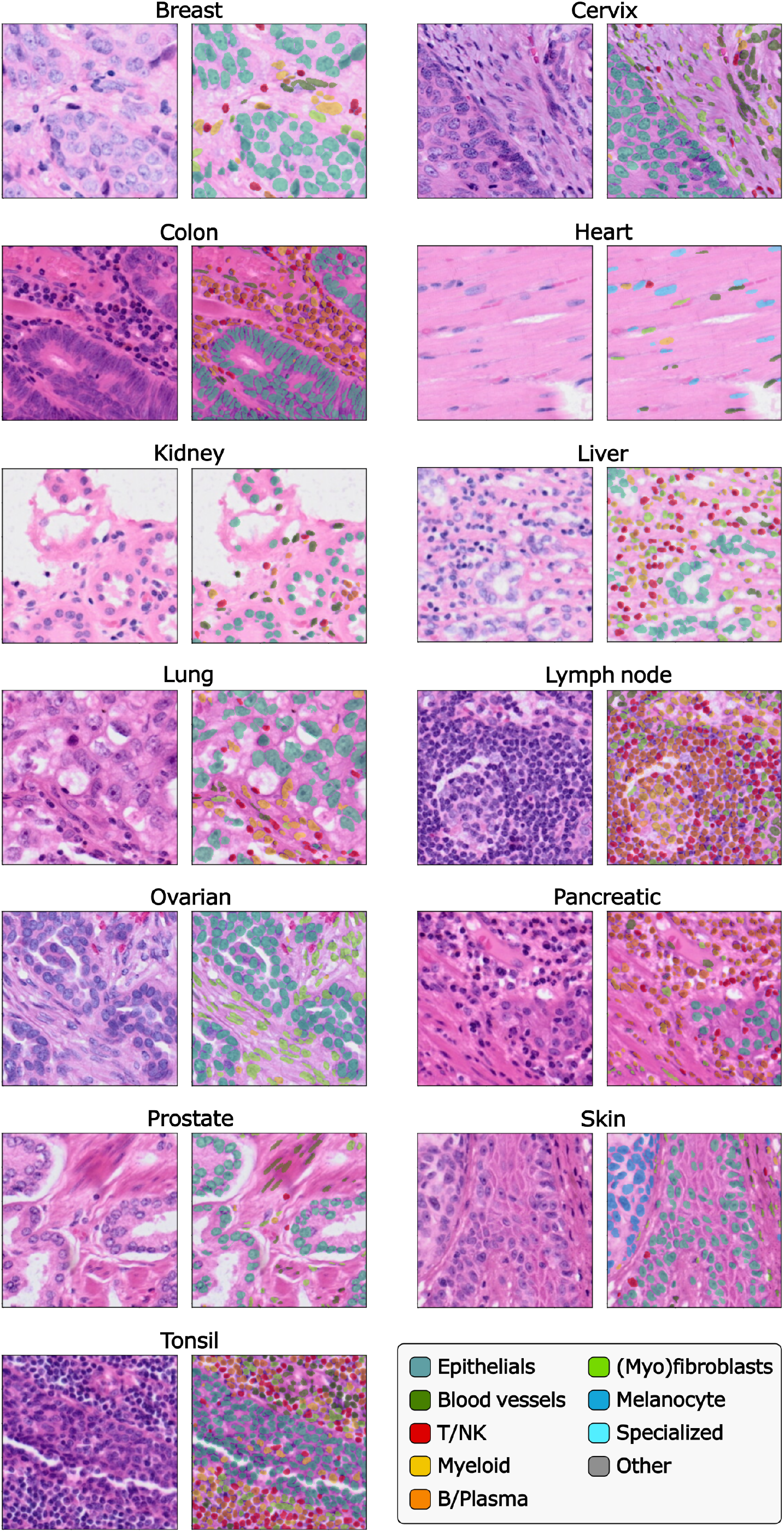
Example of 20x resolution H&E patches with corresponding segmentation and classification masks for each tissue type.

## Technical Validation

### Cell type annotation

While nucleus segmentation comes from DAPI staining for all the slides, cell segmentation from 10x Genomics is determined by two main methods, according to available data. The first, nuclear expansion method, approximates cell boundaries by isotropically expanding these nuclear boundaries until encountering adjacent cells or a predefined limit. The second, a multimodal method using additional cytoplasmic and membrane staining alongside DAPI staining, produced more accurate boundaries ^16^. Cell boundaries critically determine the RNA count attributed to each gene within individual cells, and thus inaccuracies in cell boundary delineation directly influence RNA quantification, potentially resulting in inaccurate cell annotations. Conversely, only relying on more precise nucleus boundaries risks omitting significant cytoplasmic gene expression information. To evaluate the impact of boundaries selection on cell type annotations, additional analyses were performed. First, the annotation pipeline was independently applied to two RNA count matrices: one based on nuclear boundaries and the other based on whole-cell boundaries. After filtering out cells with fewer than 10 transcripts, the concordance between annotations derived from these two matrices was 80.4% *±* 8.9. Additionally, we employed a second validation approach by applying Multiple Factor Analysis (MFA) ^63^ on cytoplasmic and nuclear RNA count data, followed again by Leiden clustering. Within each resulting cluster, we compared the proportions of annotations derived exclusively from nuclear data to those derived exclusively from whole-cell data. Clusters showing high agreement adopted labels from the whole-cell approach, while clusters with significant discrepancies underwent additional differential expression analysis to assign definitive labels. Finally, these MFA-based combined labels were compared with those previously obtained from the whole-cell analyses. This comparison showed a concordance of 83.3% ± 7.4.

Further validation of cell annotations was accomplished using the single-cell Variational Inference (scVI) model, a variational autoencoder (VAE)-based deep learning framework specialized for single-cell RNA sequencing analysis ^52^. scVI models were trained independently on each slide. UMAP visualizations of scVI embeddings, colored by our final cell labels, facilitated the qualitative assessment of annotation consistency. Results generally demonstrate strong agreement, with illustrative examples shown in Figure 7.

**Figure 7.**
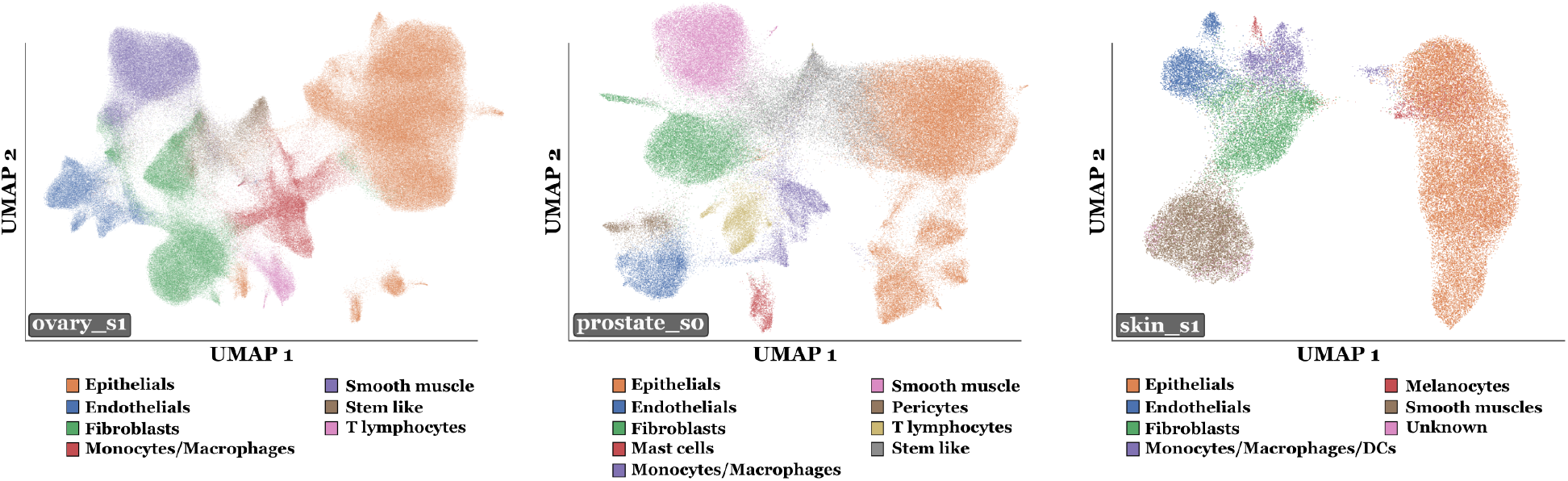
UMAPs from scVI embeddings for three example slides “ovary_s1”, “prostate_s0” and “skin_s1”. An scVI model was trained on each slide separately using the RNA count tables for the cells. UMAPs were then performed on the embeddings obtained with the trained scVI model. Labels from the previous annotation process with Leiden clustering are used.

Additionally, total RNA count values per cell were discretized into seven distinct categories and subsequently visualized via UMAP. This approach confirmed that cell clustering patterns were driven by specific gene expression profiles rather than by variations in total RNA quantity.

Finally, labels were qualitatively and globally reviewed by a pathologist, using Scverse’s napari visualization tool to simultaneously examine cell annotations and corresponding H&E images, providing expert validation of annotation accuracy and biological plausibility.

### Patching and segmentation quality

H&E image patches of size 256 × 256 px with an overlap of 64 px were generated for each slide. Using the previously established cell annotations and the spatial alignment between ST and H&E data, masks for nuclei segmentation and cell type classification were created. To evaluate the quality of segmentation and alignment, each generated mask was compared with segmentation predictions obtained from the pre-trained SAM-H/Hovernet CellViT model ^7^. The Dice coefficient, Jaccard index, and Panoptic Quality (bPQ) metrics were employed for quantitative evaluation, with the resulting distributions across all patches (“brain_s0”, “bone_marrow_s0”, “bone_marrow_s1”, “bone_s0” slides excluded) visualized in Figure 8.

**Figure 8.**
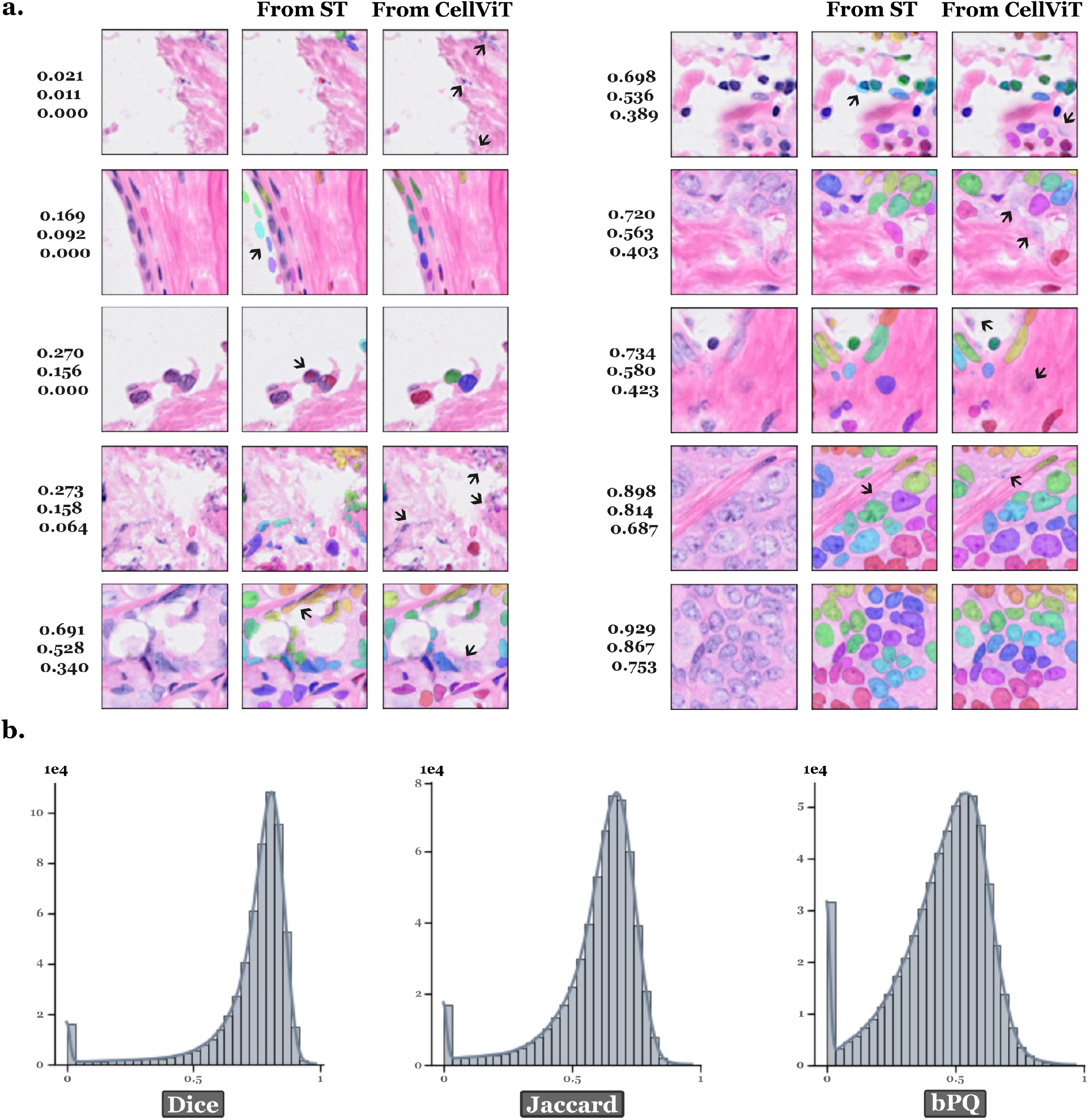
Pre-trained CellViT’s evaluation of the segmentation. The CellViT model pre-trained by the authors on the PanNuke dataset was applied to each slide. The predicted binary mask is compared with the segmentation mask coming from ST. The latter was generated using the segmentation on the DAPI and aligning the slide with the H&E modality. The comparison between the two masks for each patch is made using Dice coefficient, Jaccard index and bPQ scores. **a.** Examples of patches for “breast s0” with from left to right: H&E patch only, H&E patch with the mask from ST, and H&E patch with the mask generated by the pre-trained CellViT model. Patch scores are displayed at the left of each patch in the following order: Dice coefficient, Jaccard index and bPQ. Color identifies each unique nucleus and is not related to cell type here. Arrows highlight segmentation error examples. **b.** Distribution of Dice, Jaccard and bPQ scores for each patch over all the slides (“brain_s0”, “bone_marrow_s0”, “bone_marrow_s1”, “bone_s0” excluded).

The metric distributions demonstrate overall robust segmentation and alignment quality. To elucidate discrepancies between our segmentation and the CellViT predictions, representative patches illustrating both segmentation outputs for “breast s0” slide are presented in Figure 8. Each resulting score value corresponds to distinct scenarios. For instance, low scores may reflect regions with very few nuclei and they are undetected by one segmentation method, or significant misalignment between modalities. A low score can also correspond to poor image quality, for example due to blurring or other artifacts. Intermediate scores typically indicate minor alignment deviations or partial nuclei detection by either method. High scores generally indicate strong segmentation agreement, although minor detection discrepancies may persist. Importantly, a low segmentation score does not inherently indicate poor segmentation accuracy for ST mask; it may also arise from challenging detection scenarios specifically problematic for the CellViT model. Thus, while filtering based on segmentation metrics effectively reduces misalignment issues, it may unintentionally exclude patches containing nuclei that are inherently challenging to detect, potentially identifiable only through DAPI staining. Nonetheless, patches with higher metric scores indicate reliable segmentation quality, meaning that even though some informative data might be lost during filtering, the resulting dataset will overall have enhanced quality. These segmentation quality scores have been incorporated into the dataset to facilitate dynamic filtering, allowing flexibility in selecting patches according to specific analytical requirements.

### Dataset for fine-tuning the CellViT model

To demonstrate the utility of our dataset for downstream predictive tasks, we fine-tuned the CellViT model using STHELAR annotations. The patch dataset was processed following the CellViT authors’ data preparation pipeline, converting annotations masks into the input format required by the CellViT model. Given the large number of patches and categories, file storage formats were adapted for sparse representation to mitigate excessive storage requirements. A flexible pipeline was established to dynamically use various dataset configurations without requiring the reconstruction of data. Two distinct versions of the dataset were prepared for separate fine-tuning experiments. The first version employed detailed cell-type labels as described above, while the second adopted broader labels expected to facilitate more straightforward predictions. Detailed labels, such as distinguishing T lymphocytes from B lymphocytes, are often challenging due to their morphological similarities, which even experienced pathologists find difficult to differentiate. To get more generalized labels, specific cell categories were grouped as follows: “Immune” combined “T_NK”, “B_Plasma”, and “Myeloid”; “Stromal” merged “Blood_vessel” and “Fibroblast_Myofibroblast”; and “Other” grouped “Specialized” and the original “Other” category. Additionally, both dataset versions underwent quality filtering based on segmentation metrics, retaining only patches with a Jaccard score above 0.45. Consequently, this filtering step resulted in a selection of 508,483 patches at 40x from an original total of 587,555. The final dataset after patching and filtering is illustrated in Figure 9. Since a cell can be cut off when dividing WSIs into patches, and because there is a 64 px overlap between patches, many cells appear in multiple patches. Therefore, if we count the number of instances after patching, treating each occurrence of a cell on a patch as unique, the total number of cells increases significantly. This explains why the total number of cells is now around 19.7M, higher than the actual number of cells in the original dataset.

**Figure 9.**
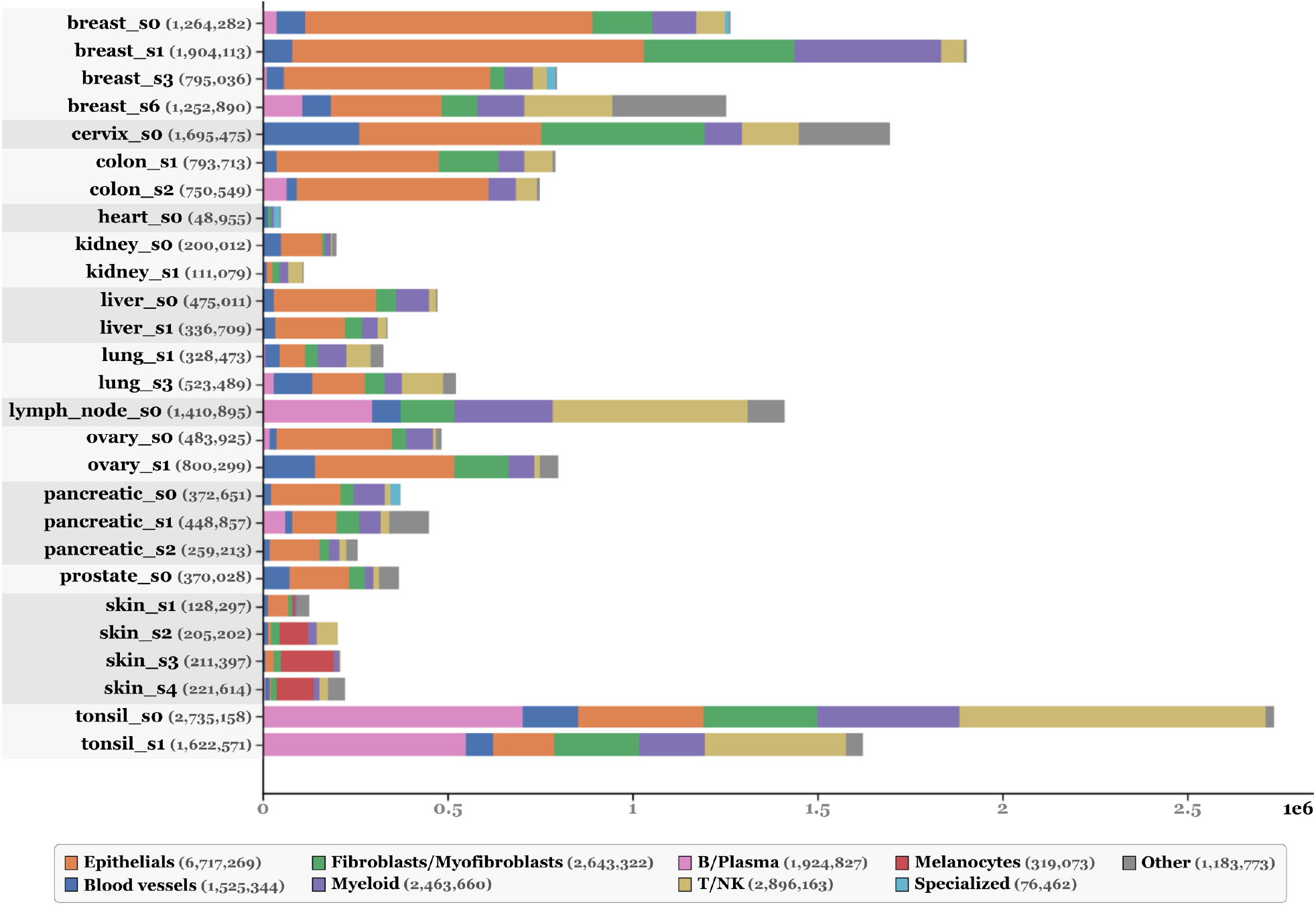
Nuclei count for each slide in the dataset after making H&E patches and after filtering patches with a Jaccard index below 0.45. The total number of nuclei for each slide or for each cell type is shown in parenthesis. The count is in logarithmic scale. The same cell can be counted several times since it can be present on several patches. Inspired from CellViT article ^7^.

### CellViT fine-tuning

The CellViT model ^67^ was fine-tuned independently for each dataset configuration. In each case, the data were split into training (60%), validation (20%), and test (20%) sets, with care taken to ensure a balanced distribution of slides and cell types across all subsets.

Fine-tuning was performed for the CellViT model variant using a Segment Anything Model (SAM)-based encoder and a HoVer-Net-style decoder. The model architecture remained unchanged, except for the NT branch, where the final decoder layer was adapted to accommodate our custom set of cell type labels. Initialization was conducted using pre-trained weights from the CellViT-SAM-H model 40x, which employs a ViT-Huge encoder with 632 million parameters. This encoder includes 32 transformer blocks, 16 attention heads, and an embedding dimension of 1280.

The final fine-tuning did not use the authors’ original sampling strategy, as preliminary tests indicated that the class-weight strategy applied to the focal Tversky (FT) loss for the NT branch was sufficient. The loss weighting scheme used during training was consistent with the original configuration. Specifically, the TC branch used a weight of *λ* = 0.1 for the cross-entropy loss. The NP branch applied *λ* = 1.0 for both the Focal Tversky loss and Dice loss. The HV branch used *λ* = 2.5 for mean squared error (MSE) and *λ* = 8.0 for mean squared gradient error (MSGE). For the NT branch, *λ* = 0.5 was used for binary cross-entropy (BCE), *λ* = 0.2 for Dice loss, and *λ* = 0.5 for the multi-class Focal Tversky loss. Within the multi-class Focal Tversky loss, class weights were adjusted using the formula *w*_*c*_ = log(1 + 1*/freq_c_*), where *freq*_*c*_ denotes the frequency of class *c* in the training set. The background weight was set equal to that of the most common cell type. As the “Other” category grouped cells with low RNA content or no clear marker genes, its contribution to the NT branch losses was downweighted to 0.1 for BCE and Dice loss, and to 0.1 times the weight of the most prevalent class for the Focal Tversky loss.

Training was performed for 53 or 42 epochs for detailed and grouped labels respectively, using mixed precision and a batch size of 32. The AdamW optimizer was employed with a learning rate of 3.10^*−*4^, a weight decay of 1.10^*−*4^, and *β* values of (0.85, 0.95). An exponential learning rate scheduler with a decay factor of *γ* = 0.85 was applied. Full model fine-tuning began from epoch 2 by unfreezing the encoder. Validation metrics were monitored throughout training, and the best models were selected based on these metrics: epoch 40 for the detailed label configuration and epoch 32 for the grouped label configuration.

Despite the large number of cells, the current dataset originates from a limited number of slides. This constraint poses a challenge for the development of generalizable models, particularly when applied to WSIs coming from independent sources or different experimental conditions. To partially mitigate this limitation, the default data augmentation pipeline was retained during training, incorporating random rotations, flips, elastic deformations, blurring, Gaussian noise, and cropping. The only modification was the replacement of color jitter augmentation with a histology-specific HED color augmentation ^64^. All images were normalized using mean and standard deviation values of (0.5, 0.5, 0.5).

Performance results are summarized in Table 3 for detailed labels, and Table 4 for grouped labels. These results indicate that the CellViT model, originally trained on the PanNuke dataset, can effectively adapt to our Xenium-derived dataset through targeted fine-tuning. When trained on the nine detailed cell-type classes, the model retains excellent binary segmentation performance inherited from the pre-training (mean Dice = 0.883, mean Jaccard = 0.794, F1 = 0.855, and bPQ = 0.652). Despite nearly doubling the complexity of the label space and introducing morphologically challenging immune cell subtypes, the class-aware Panoptic Quality (mPQ) remains respectable at 0.369 (compared to 0.496 for initial CellViT performance on PanNuke). Performance is notably strong for morphologically distinct phenotypes, with Panoptic Quality (PQ) scores of 0.541 and 0.535 and F1 scores of 0.720 and 0.772 for epithelial cells and melanocytes, respectively. Classification is more challenging for inflammatory cell subtypes, yet performance remains encouraging, with PQ values ranging from 0.208 to 0.336 and F1 scores between 0.329 and 0.428. For the different connective cell types ((myo)fibroblast cells and blood vessel), PQ and F1 scores were 0.343 and between 0.404 and 0.490, respectively. To compare with pre-trained CellViT on PanNuke, the global inflammatory and connective categories achieves PQ values of 0.417 and 0.423, and F1 scores of 0.58 and 0.53, respectively. Cells categorized as “Other” have relatively low scores, which is expected given the intentionally reduced training weight assigned to this category. This decision reflects the inherent ambiguity and limited RNA content of these cells, allowing the model greater flexibility in classification. Segmentation quality is consistent across various tissues, indicated by Dice scores between 0.847 and 0.908 and bPQ values from 0.539 to 0.727.

**Table 3:** Evaluation on the test set using CellViT fine-tuned with detailed cell type annotations.

| Binary dataset metrics |  |  |  |  |
| --- | --- | --- | --- | --- |
| Binary-Cell-Dice-Mean |  |  |  | 0.883 |
| Binary-Cell-Jaccard-Mean |  |  |  | 0.794 |
| Tissue-Multiclass-Accuracy |  |  |  | 0.999 |
| bPQ / bDQ / bSQ |  | 0.652 / 0.820 / 0.792 |  |  |
| mPQ / mDQ / mSQ |  | 0.369 / 0.462 / 0.572 |  |  |
| F1 / precision / recall (detection) |  | 0.855 / 0.884 / 0.829 |  |  |
| Tissue-specific metrics |  |  |  |  |
| Tissue | Dice | Jaccard | mPQ | bPQ |
| Breast | 0.895 | 0.812 | 0.392 | 0.665 |
| Cervix | 0.889 | 0.802 | 0.375 | 0.671 |
| Colon | 0.847 | 0.737 | 0.316 | 0.539 |
| Heart | 0.867 | 0.768 | 0.317 | 0.644 |
| Kidney | 0.905 | 0.828 | 0.390 | 0.727 |
| Liver | 0.908 | 0.833 | 0.444 | 0.725 |
| Lung | 0.877 | 0.783 | 0.352 | 0.652 |
| LymphNode | 0.840 | 0.726 | 0.262 | 0.609 |
| Ovarian | 0.879 | 0.786 | 0.384 | 0.656 |
| Pancreatic | 0.863 | 0.762 | 0.335 | 0.596 |
| Prostate | 0.859 | 0.755 | 0.378 | 0.630 |
| Skin | 0.874 | 0.780 | 0.390 | 0.640 |
| Tonsil | 0.865 | 0.763 | 0.257 | 0.630 |
| Nuclei type metrics |  |  |  |  |
| Nuclei Type | DQ | SQ | PQ |  |
| T_NK | 0.409 | 0.544 | 0.336 |  |
| B_Plasma | 0.259 | 0.450 | 0.208 |  |
| Myeloid | 0.370 | 0.547 | 0.294 |  |
| Blood_vessel | 0.441 | 0.537 | 0.343 |  |
| Fibroblast_Myofibroblast | 0.438 | 0.595 | 0.343 |  |
| Epithelial | 0.664 | 0.732 | 0.541 |  |
| Specialized | 0.242 | 0.300 | 0.195 |  |
| Melanocyte | 0.652 | 0.703 | 0.535 |  |
| Other | 0.228 | 0.346 | 0.170 |  |
| Nuclei detection metrics |  |  |  |  |
| Nuclei Type | Precision | Recall | F1 |  |
| T_NK | 0.401 | 0.460 | 0.428 |  |
| B_Plasma | 0.428 | 0.365 | 0.394 |  |
| Myeloid | 0.367 | 0.298 | 0.329 |  |
| Blood_vessel | 0.559 | 0.437 | 0.490 |  |
| Fibroblast_Myofibroblast | 0.422 | 0.387 | 0.404 |  |
| Epithelial | 0.705 | 0.736 | 0.720 |  |
| Specialized | 0.580 | 0.282 | 0.379 |  |
| Melanocyte | 0.791 | 0.755 | 0.772 |  |
| Other | 0.453 | 0.305 | 0.365 |  |

**Table 4:**
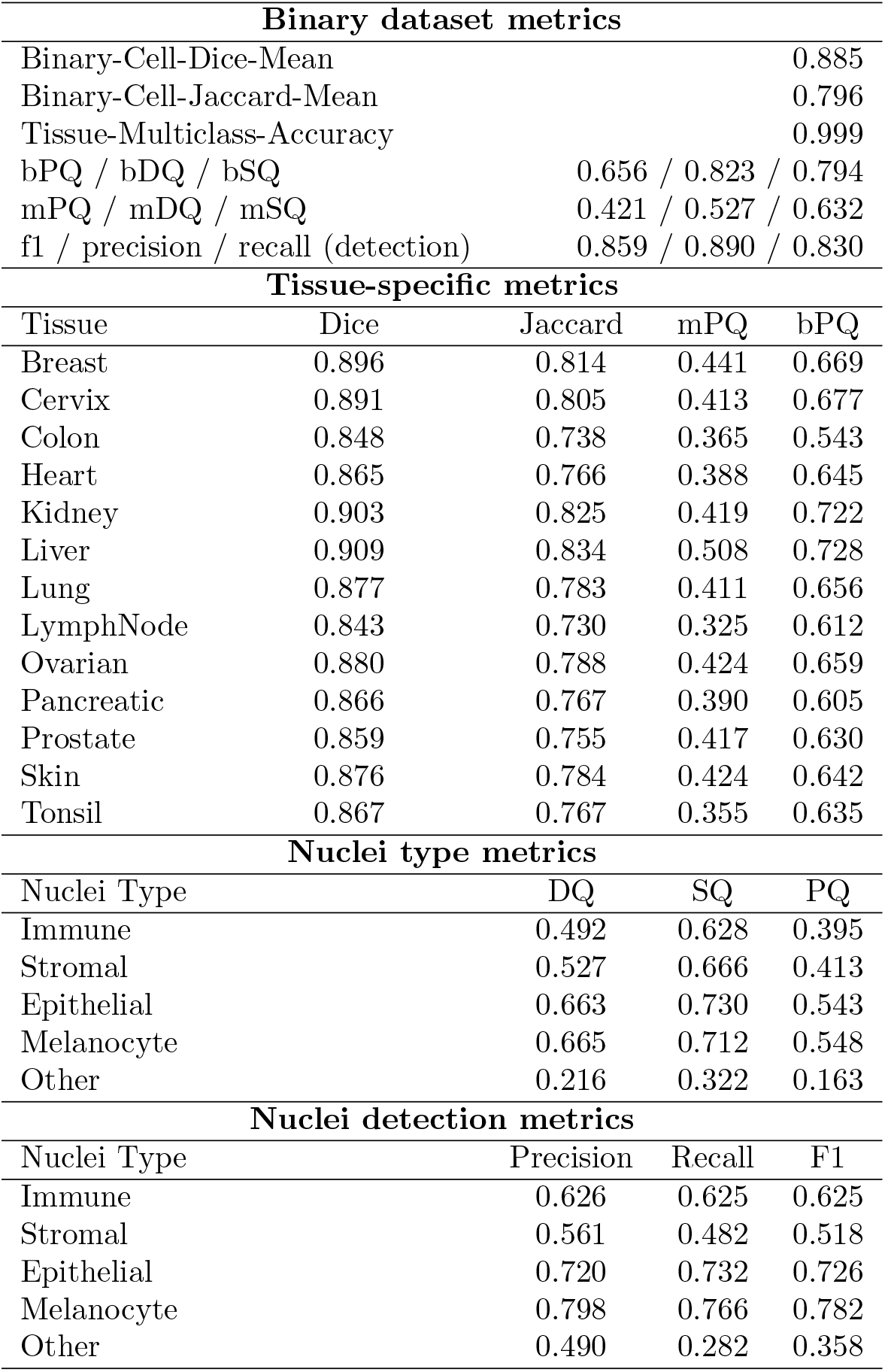
Evaluation on the test set using CellViT fine-tuned with grouped cell type annotations.

| Binary dataset metrics |  |  |  |  |
| --- | --- | --- | --- | --- |
| Binary-Cell-Dice-Mean |  |  | 0.885 |  |
| Binary-Cell-Jaccard-Mean |  |  | 0.796 |  |
| Tissue-Multiclass-Accuracy |  |  | 0.999 |  |
| bPQ / bDQ / bSQ |  |  | 0.656 / 0.823 / 0.794 |  |
| mPQ / mDQ / mSQ |  |  | 0.421 / 0.527 / 0.632 |  |
| f1 / precision / recall (detection) |  |  | 0.859 / 0.890 / 0.830 |  |
| Tissue-specific metrics |  |  |  |  |
| Tissue | Dice | Jaccard | mPQ | bPQ |
| Breast | 0.896 | 0.814 | 0.441 | 0.669 |
| Cervix | 0.891 | 0.805 | 0.413 | 0.677 |
| Colon | 0.848 | 0.738 | 0.365 | 0.543 |
| Heart | 0.865 | 0.766 | 0.388 | 0.645 |
| Kidney | 0.903 | 0.825 | 0.419 | 0.722 |
| Liver | 0.909 | 0.834 | 0.508 | 0.728 |
| Lung | 0.877 | 0.783 | 0.411 | 0.656 |
| LymphNode | 0.843 | 0.730 | 0.325 | 0.612 |
| Ovarian | 0.880 | 0.788 | 0.424 | 0.659 |
| Pancreatic | 0.866 | 0.767 | 0.390 | 0.605 |
| Prostate | 0.859 | 0.755 | 0.417 | 0.630 |
| Skin | 0.876 | 0.784 | 0.424 | 0.642 |
| Tonsil | 0.867 | 0.767 | 0.355 | 0.635 |
| Nuclei type metrics |  |  |  |  |
| Nuclei Type |  | DQ | SQ | PQ |
| Immune |  | 0.492 | 0.628 | 0.395 |
| Stromal |  | 0.527 | 0.666 | 0.413 |
| Epithelial |  | 0.663 | 0.730 | 0.543 |
| Melanocyte |  | 0.665 | 0.712 | 0.548 |
| Other |  | 0.216 | 0.322 | 0.163 |
| Nuclei detection metrics |  |  |  |  |
| Nuclei Type |  | Precision | Recall | F1 |
| Immune |  | 0.626 | 0.625 | 0.625 |
| Stromal |  | 0.561 | 0.482 | 0.518 |
| Epithelial |  | 0.720 | 0.732 | 0.726 |
| Melanocyte |  | 0.798 | 0.766 | 0.782 |
| Other |  | 0.490 | 0.282 | 0.358 |

When closely related phenotypes, difficult to distinguish based only on H&E morphology, were merged into five broader classes, classification metrics improved significantly without compromising segmentation performance. In comparison with the pre-trained CellViT results, the fine-tuned model achieved PQ scores of 0.395 for immune cells (versus 0.417 for inflammatory for pre-trained), 0.413 for stromal cells (versus 0.423 for connective for pre-trained), and 0.543 and 0.548 for epithelial and melanocyte categories (versus 0.581 and 0.583 for neoplastic and epithelial in the pre-trained case). Correspondingly, F1 scores improved to 0.625 for immune (versus 0.58 for inflammatory for pre-trained), 0.518 for stromal (versus 0.53 for connective for pre-trained), and 0.726 and 0.782 for epithelial and melanocyte categories (versus 0.71 and 0.73 for neoplastic and epithelial in the pre-trained case).

These performance metrics, particularly PQ, can be impacted by the accuracy of segmentation. Given that our ground-truth segmentation may exhibit minor pixel-level inaccuracies due to slight misalignments, we further investigated model performance using a confusion matrix within the matched pairs prediction/ground-truth nuclei obtained through the Kuhn-Munkres algorithm (Table 5). This analysis confirms overall agreement. As expected, it is more difficult to differentiate B and T lymphocytes that are nearly indistinguishable morphologically in H&E. Future improvements might be to restrict the test set to patches with the highest segmentation quality scores. This approach could ensure greater confidence in performance metrics by minimizing the confounding effects of segmentation inaccuracies.

**Table 5:**
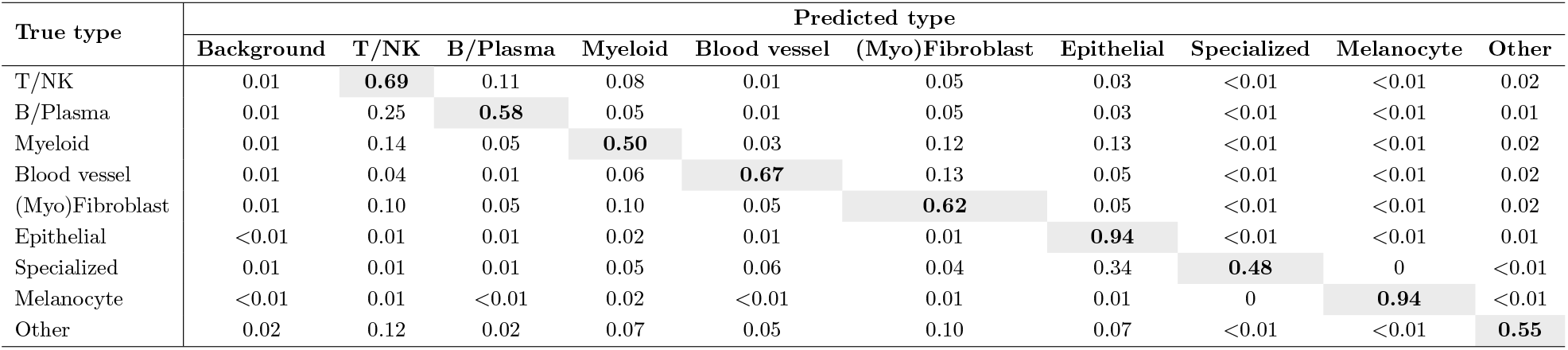
Cell type classification confusion matrix on the test set. The confusion matrix compares ground truth labels and prediction from the CellViT model fine-tuned on detailed cell types. The comparison was done only within the matched pairs ground truch/predicted nuclei obtained through the Kuhn-Munkres algorithm.

| True type | Predicted type |  |  |  |  |  |  |  |  |  |
| --- | --- | --- | --- | --- | --- | --- | --- | --- | --- | --- |
|  | Background | T/NK | B/Plasma | Myeloid | Blood vessel | (Myo)Fibroblast | Epithelial | Specialized | Melanocyte | Other |
| T/NK | 0.01 | **0.69** | 0.11 | 0.08 | 0.01 | 0.05 | 0.03 | <0.01 | <0.01 | 0.02 |
| B/Plasma | 0.01 | 0.25 | **0.58** | 0.05 | 0.01 | 0.05 | 0.03 | <0.01 | <0.01 | 0.01 |
| Myeloid | 0.01 | 0.14 | 0.05 | **0.50** | 0.03 | 0.12 | 0.13 | <0.01 | <0.01 | 0.02 |
| Blood vessel | 0.01 | 0.04 | 0.01 | 0.06 | **0.67** | 0.13 | 0.05 | <0.01 | <0.01 | 0.02 |
| (Myo)Fibroblast | 0.01 | 0.10 | 0.05 | 0.10 | 0.05 | **0.62** | 0.05 | <0.01 | <0.01 | 0.02 |
| Epithelial | <0.01 | 0.01 | 0.01 | 0.02 | 0.01 | 0.01 | **0.94** | <0.01 | <0.01 | 0.01 |
| Specialized | 0.01 | 0.01 | 0.01 | 0.05 | 0.06 | 0.04 | 0.34 | **0.48** | 0 | <0.01 |
| Melanocyte | <0.01 | 0.01 | <0.01 | 0.02 | <0.01 | 0.01 | 0.01 | 0 | **0.94** | <0.01 |
| Other | 0.02 | 0.12 | 0.02 | 0.07 | 0.05 | 0.10 | 0.07 | <0.01 | <0.01 | **0.55** |

An increase in the number of available slides would also make it possible to improve the overall segmentation quality of the dataset. With a larger dataset, we can apply more stringent filtering criteria during patch selection, retaining only regions with excellent segmentation and accurate alignment between modalities. This selective approach would raise the average quality of the final dataset. In parallel, improving the criteria used for patch selection may further enhance consistency. For instance, switching to a selection using PQ metric, which simultaneously accounts for segmentation and classification performance, could provide a more comprehensive and quantitative basis for patch evaluation. Additionally, using more advanced alignment tools, such as the Valis method, could help correct residual spatial misalignments in certain patches. A combined approach that leverages stricter selection, better metrics, and improved alignment techniques may offer the most robust solution. Interestingly, since the pre-trained CellViT’s predictions have been computed for each patch, the dataset may also be used to evaluate modality realignment methods, highlighting an additional potential application.

### PanNuke labels

To assess the cell type prediction performance of pre-trained CellViT, the model was applied to our dataset in inference mode. The architecture used was consistently the SAM-H/HoVer-Net variant of CellViT, pretrained on PanNuke. To ensure a direct comparison between predicted cell types and those obtained through RNA-based clustering, predicted labels from CellViT output were not used directly. Instead, pixel-level probability maps for each cell type were retrieved from the NT branch output, and the post-processing was implemented using using Xenium-derived segmentation masks. Multiple evaluation strategies were applied, including confusion matrices, Sankey diagrams, and UMAP visualizations. These analyses were performed at both the full dataset level and on a slide-by-slide basis to explore prediction consistency and discrepancies. The same analysis pipeline was applied to the outputs of our fine-tuned CellViT models, primarily for exploratory purposes. These comparisons offer insight into persistent sources of confusion between cell types, but interpretation remains limited by the fact that cells are not coming only from the evaluation set but also from training and validation sets.

Table 6 provides an overview of the results for the pre-trained CellViT model. Myeloid cells seems to be particularly challenging to predict, as they are frequently misclassified as connective instead of inflammatory. Furthermore, contrary to expectations, cells with fewer than 10 RNAs are rarely classified as dead cells, potentially highlighting a limitation in either the prediction capability or the underlying assumption that low RNA content directly indicates cell death. Distinguishing between normal epithelial and neoplastic cells seems also problematic. While a predominant classification of epithelial cells as neoplastic is reasonable considering the number of cancerous tissues, some normal epithelial cells should be still expected. For instance, the slide “liver s0” is derived from a healthy patient, but it exhibits a 64% classification rate of epithelial cells as neoplastic, highlighting this limitation. It should also be mentioned that heart, tonsil, and lymph node tissues are present in our dataset but absent from the original PanNuke dataset, potentially affecting model generalization and accuracy for these specific tissues.

**Table 6:** Comparison between RNA-based cell type annotations and PanNuke labels predicted by the pre-trained CellViT model (row-normalized). Each row corresponds to a cell type defined using transcript information. Columns indicate the proportion of cells assigned to each PanNuke category by the CellViT model, using Xenium-derived segmentation masks.

| Cell type | PanNuke label |  |  |  |  |  |
| --- | --- | --- | --- | --- | --- | --- |
|  | Background | Connective | Dead | Epithelial | Inflammatory | Neoplastic |
| B_Plasma | 0.05 | 0.12 | 0 | 0.01 | **0.60** | 0.21 |
| Blood_vessel | 0.12 | **0.68** | 0 | 0.01 | 0.11 | 0.09 |
| Epithelial | 0.10 | 0.24 | 0 | 0.03 | 0.03 | **0.60** |
| Fibroblast_Myofibroblast | 0.12 | **0.62** | 0 | 0.01 | 0.16 | 0.09 |
| Less10 | 0.30 | **0.42** | 0.01 | 0.01 | 0.18 | 0.07 |
| Melanocyte | 0.12 | 0.11 | 0 | 0.02 | 0.01 | **0.75** |
| Myeloid | 0.11 | **0.47** | 0 | 0.01 | 0.23 | 0.18 |
| Specialized | 0.10 | **0.56** | 0 | 0.01 | 0.04 | 0.29 |
| Stem_like | 0.18 | **0.56** | 0 | 0.01 | 0.09 | 0.16 |
| T_NK | 0.04 | 0.20 | 0 | 0.01 | **0.60** | 0.15 |
| Unknown | 0.19 | **0.46** | 0.01 | 0.01 | 0.23 | 0.09 |

Overall, we can see that the broad categorization employed by PanNuke inherently results in loss of detailed cellular information. Hence, exploring approaches for more refined predictions appears promising for better characterizing cell types.

### H&E features

To compare cell-level embeddings derived from H&E images with those obtained from RNA data, three different encoders were applied to the H&E patches for the whole dataset. In addition to the pre-trained CellViT encoder, two other pre-trained ViT were evaluated. The first was the Phikon-v2 model, developed by Owkin, which is a ViT Large pre-trained using the DinoV2 self-supervised learning approach on the PANCAN-XL dataset ^65^. PANCAN-XL comprises approximately 450 million histology image tiles at 20x magnification, sampled from over 60,000 whole slide images. The second model was Google’s ViT-Base-Patch16-224-in21k, a transformer encoder (BERT-like architecture) trained in a supervised manner on the ImageNet-21k dataset ^66^. For all three models, cell-level embeddings were extracted as follows: for each cell in a given patch, sub-patches containing at least one pixel belonging to that cell were identified. In this context, “sub-patches” refer to the input tokens used by transformer architectures. Embeddings were extracted for each relevant sub-patch and then averaged to produce a single patch-level embedding per cell. When a cell appeared in multiple patches, the embeddings from those patches were further averaged to generate a single global embedding per cell. For CellViT, nucleus segmentation was used to define relevant sub-patches, while for the Phikon-v2 and Google models, cell segmentation boundaries were used. For each model and each slide, PCA was performed on the resulting cell embeddings. We retained the minimum number of principal components required to explain 95% of the total variance, and used these components as input for k-means clustering. The optimal number of clusters was determined using the Elbow method combined with the KneeLocator algorithm. Cluster assignments were visualized on the UMAP projections of the RNA-derived data, allowing for direct comparison of clustering patterns between the two modalities. This whole process was repeated using then the fine-tuned CellViT models, to assess whether fine-tuning improved alignment between H&E-derived and RNA-derived embeddings, particularly for models trained with more refined cell type labels. However, as with earlier comparisons, it is important to interpret these results cautiously, as the features were extracted from all cells and not exclusively from the independent test set.

Representative examples of feature evolution are shown in Figure 10, where RNA-based embeddings are projected using UMAP and colored by k-means cluster assignments derived from each H&E encoder. Visual alignment is considered better when these clusters correspond to well-separated regions in the RNA-derived space. Observations indicate that fine-tuning CellViT improved the alignment between H&E-derived clusters and RNA-derived clusters for some slides. This improvement is not consistently observed across all slides, which is not surprising given the diversity and complexity of the data. Moreover, improved clustering alignment does not necessarily predict enhanced performance in subsequent tasks, although these results remain insightful. The alignment is also superior to that observed with both Phikon-v2 and Google’s ViT.

**Figure 10.**
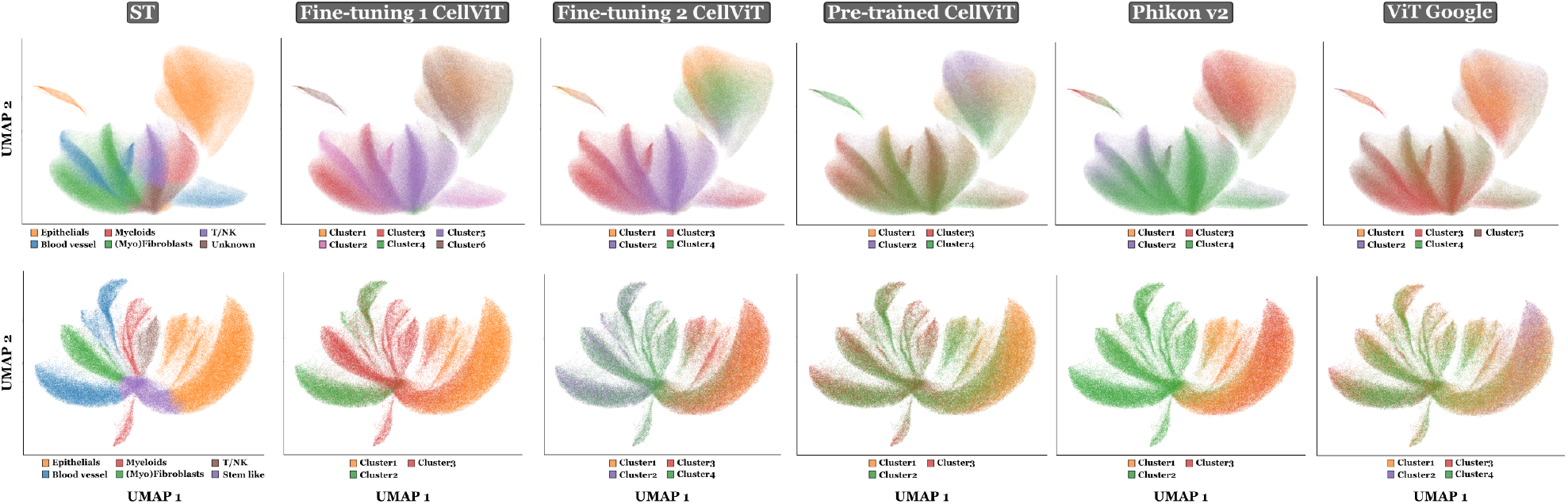
Comparison of H&E-derived features for slides “cervix_s0” (top) and “prostate_s0” (bottom). UMAPs are based on RNA-derived embeddings used during the cell type annotation process. Cluster colors (left to right) represent: (1) cell types assigned from RNA-based labeling, k-means clusters derived from nucleus features extracted using (2) CellViT fine-tuned on detailed cell types, (3) CellViT finetuned on grouped cell types, (4) pre-trained CellViT, and k-means clusters derived from cell features extracted using (5) Phikon-v2 encoder, and (6) ViT-Base encoder from Google.

An alternative approach to mitigate the dependency on segmentation quality would be to use the dataset on the recently proposed CellViT++ ^67^. In this pipeline, CellViT is first applied to extract both segmentation masks and feature embeddings for each nucleus. These embeddings can then be used as input for training a separate classifier, allowing the prediction of other cell types. This strategy offers a promising path for leveraging the dataset’s biological content, particularly in cases where segmentation may not be perfectly aligned between modalities, even if a baseline level of segmentation quality remains essential to ensure correspondence between the segmentation masks in our dataset and those produced by CellViT. Nevertheless, this decoupling has limitations since the CellViT++ solution does not enable to fine-tune the encoder. As a result, if the pre-trained features lack the discriminative information required to distinguish between specific cell types, their classification by the classifier will be limited. Our experiments comparing nucleus-level features using k-means clustering before and after CellViT encoder fine-tuning suggest that adjusting the feature space could be relevant for more precise cell type classification.

## Usage Notes

Xenium data from 10x Genomics dataset platform is licensed under the Creative Commons Attribution 4.0 International (CC BY 4.0) license.

## Code Availability

Source code is available at on github with the two following repositories:

### STHELAR

Code for generating the dataset and validation analysis, accessible at https://github.com/MICS-Lab/STHELAR.

### STHELAR_for_CellViT

Code from the CellViT repository ^68^ and adapted for our dataset, accessible at https://github.com/MICS-Lab/CellViT_for_STHELAR.

## Author Contributions

QB, SC, PHC designed the study. FGS conducted the study, developed the source code, developed the annotation method, ran all the tests, published the data and library, wrote the manuscript. AM is the pathologist who performed the visual qualitative annotation. QB, HB, SC provided technical support. PHC, SC supervised the study. All authors reviewed the paper.

## Acknowledgements

This work was granted access to the HPC resources of IDRIS under the allocation 2025-AD011015999 made by GENCI.

This work has benefited from the ANR grant under the investment program France 2030: PRISM ANR-23-IAHU-0002.

## Annexes

### Overview of the CellViT model

The CellViT architecture has been described in the original publication ^7^. In summary, this model consists of a Vision Transformer (ViT) encoder connected via skip connections to four separate decoder modules. These decoders are organized into three distinct multitask output branches, each performing specific segmentation tasks. The first branch predicts binary segmentation maps for all nuclei (NP), delineating their precise boundaries and shapes. The second branch generates horizontal and vertical distance maps (HV), facilitating accurate localization and segmentation of individual nuclei. The third branch predicts nuclei type maps (NT) for classifying cells into distinct types. Postprocessing involves separating overlapping nuclei and assigning nuclei types through a majority voting mechanism applied within each nuclear region, based on the predicted NT map. Additionally, a tissue classification (TC) branch is implemented ^7^.

The total loss function used to train the CellViT model is defined as:

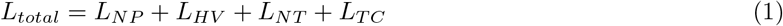

where *L*_*NP*_ denotes the loss for the NP-branch, *L*_*HV*_ the loss for the HV-branch, *L*_*NT*_ the loss for the NT-branch, and *L*_*T C*_ the loss for the TC-branch. Each individual branch loss is formulated as a weighted combination of specific loss functions:

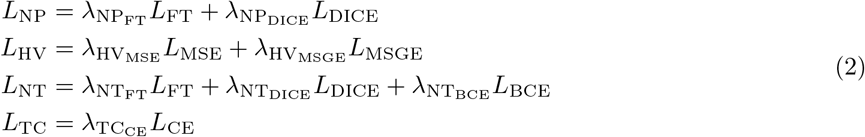

with *L*_*CE*_ the cross-entropy loss, *L*_*BCE*_ the binary cross-entropy loss, *L*_*DICE*_ the Dice loss, *L*_*F T*_ the Focal Tversky loss, *L*_*MSGE*_ the mean squared error of the gradients, and *L*_*MSE*_ the mean squared error.

